# Commonalities and differences in the Oxidative Phosphorylation of Mitochondria and Neuronal Membranes

**DOI:** 10.1101/2020.02.18.953729

**Authors:** Silvia Ravera, Martina Bartolucci, Daniela Calzia, Alessandro M. Morelli, Isabella Panfoli

**Affiliations:** Department of Experimental Medicine, University of Genoa, Via de Toni 14, 16132 Genoa, Italy; Laboratory of Mass Spectrometry - Core Facilities, Istituto Giannina Gaslini, Via G. Gaslini 5, 16147 Genoa, Italy; Department of Pharmacy, Biochemistry Lab., University of Genoa, Viale Benedetto XV 3, 16132 Genoa, Italy

**Keywords:** ATP synthase, extramitochondrial energy production, neuronal mitochondria, oxidative phosphorylation, proton gradient accumulation

## Abstract

Mitochondria are considered the exclusive site of aerobic metabolism. However, in recent years, the functional expression of the oxidative phosphorylation (OxPhos) machinery has been reported in several other membranous structures, including the plasma membrane, endoplasmic reticulum, nucleus, myelin sheath and disks of rod outer segments. Thus, to underline commonalities and differences between extra-mitochondrial and mitochondrial aerobic metabolism, we characterized the aerobic ATP synthesis in isolated myelin sheath (IM) and rod outer segment (OS) disks, using mitochondria-enriched fractions, as a positive control. Oxygen consumption and ATP synthesis were evaluated in the presence of conventional (pyruvate + malate or succinate) and unconventional (NADH) substrates. ATP synthesis was also assayed in the presence of 10-100 µM ATP in the assay medium. Data show that IM and OS disks consumed oxygen and synthesized ATP both in the presence of conventional and unconventional respiratory substrates, while the mitochondria-enriched fraction did not utilize NADH. Only in mitochondria, ATP synthesis was progressively lost in the presence of increasing ATP concentrations. Conversely, only myelin sheath and rod OS disks produced ATP at a later time or after the removal of respiratory substrates, reflecting their ability to accumulate energy and this opens up exciting perspectives in the study of sleep. Thus, these data suggest that the extramitochondrial OxPhos in IM and rod OS displays a different behavior concerning the classic mitochondrial aerobic metabolism, representing a possible basic molecular process involved in the physiology of the nervous system.

**Significance Statement:** Mitochondria are considered the cell powerhouse, being the site of the oxidative phosphorylation (OxPhos), which produces the major part of cellular chemical energy by oxygen consumption. However, proteomics, microscopy, and biochemical analyses have described the ectopic functional expression of the OxPhos machinery also in other membranous structures, such as isolated myelin (IM) and rod outer segments (OS). The results reported in this work shows that, although the proteins involved in IM and rod OS OxPhos appear the same expressed in mitochondria, the comparison of mitochondrial and extramitochondrial OxPhos display some differences, opening a new scenario about the energy metabolism modulation.

**Graphical Abstract:** 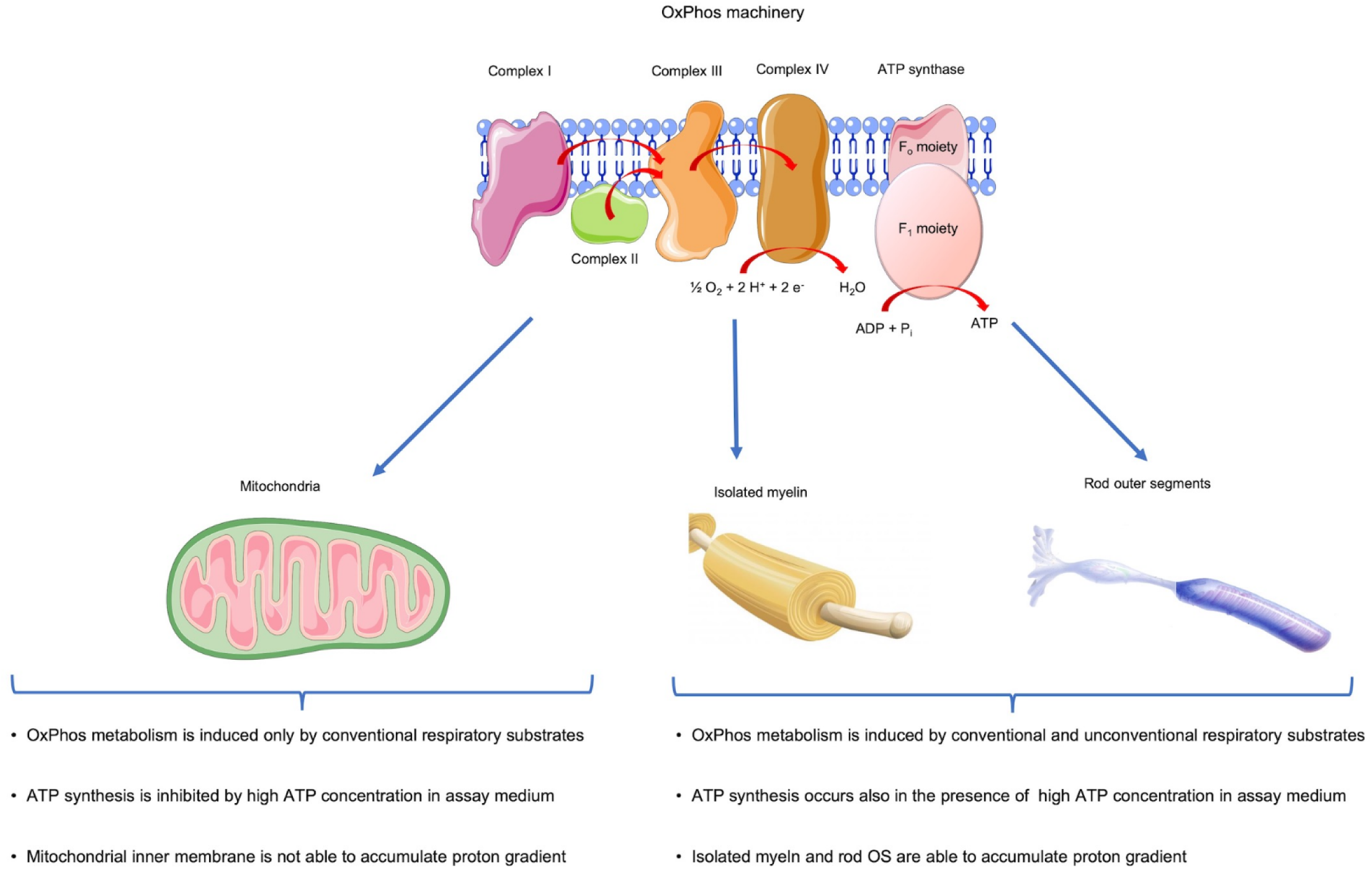

## INTRODUCTION

The aerobic metabolism is the main source of chemical energy, in terms of produced ATP (Boyer 1997), and mitochondria are considered the unique site of aerobic ATP production (Boyer 2002). However, in recent years, several biochemical and proteomic data, as well as immunofluorescence studies, demonstrated that mitochondrial components are expressed also in extra-mitochondrial membranes, specifically on the cell surface and in many subcellular membrane fractions (Bartoli *et al*. 1975; Panfoli *et al*. 2011b). In particular, the ectopic expression of F_o_-F_1_ ATP synthase, and sometimes also of the Electron Transfer Chain (ETC), has been described on plasma membrane of several cancer cells (Jung *et al*. 2007; Stockwin *et al*. 2006; Chang *et al*. 2012; Yamamoto *et al*. 2007; Young 2010) and of all of the membrane raft proteomic analyses (Panfoli *et al*. 2011b). An aerobic synthesis of ATP in cell nuclei, known to be free of mitochondria, has long been demonstrated (Klouwen and Appelman 1967). A functional ATP synthase was found on the plasma membrane of human umbilical vein endothelial cells (HUVECs) (Arakaki *et al*. 2007) and of hepatocytes (Mangiullo *et al*. 2008), as well as in isolated myelin (IM) (Ravera *et al*. 2009; Ravera *et al*. 2011; Ravera *et al*. 2013), rod OS disks (Panfoli *et al*. 2008; Panfoli *et al*. 2009; Calzia *et al*. 2013b), exosomes and microvesicles from human umbilical cord mesenchymal stem cells (Panfoli *et al*. 2016)(Bruschi *et al*. 2018) in platelets (Ravera *et al*. 2018b), in which the extramitochondrial ATP synthase is associated with the activity of ETC. Moreover, in IM and rod OS disks, the functional presence of the Krebs cycle pathway has been demonstrated (Ravera *et al*. 2013; Panfoli *et al*. 2011a), suggesting that these subcellular districts perform the whole glucose oxidation. Interestingly, the ectopic OxPhos appears composed of the same proteins expressed in mitochondria, some of which codified by the mitochondria DNA (Panfoli *et al*. 2009; Ravera *et al*. 2009). However, despite this structural identity, the functional characteristics of the ectopic oxidative metabolism seem slightly different from those of mitochondria. Therefore, in this work, to better characterize the ectopic OxPhos metabolism, we have investigated the commonalities and differences between extra-mitochondrial and mitochondrial aerobic metabolism, using IM and OS rod as models.

## METHODS AND MATERIALS

### Reagents

Salts, substrates, and all other chemicals of analytical grade were purchased from Sigma-Aldrich/Merck (St. Louis, MO, USA). For the luminometric evaluation of ATP synthesis, the ATP Bioluminescence Assay Kit CLS II, by Roche (Basel, Switzerland) was used.

Ultrapure water (Milli-Q; Millipore, Billerica, MA, USA) was used throughout. Safety precautions were taken for chemical hazards in carrying out the experiments.

### Samples

Bovine forebrain and retina of cattle of less than 1 year of age were collected in a slaughterhouse, following the safety precautions.

### Mitochondria-enriched fraction isolation

Mitochondria-enriched fraction (MIT) was obtained according to Lescuyer et al (Lescuyer *et al*. 2003). Briefly, bovine forebrain washed in PBS, was homogenized in Buffer A, containing 0.25 M sucrose, 0.15 M KCl, 10 mM TRIS HCl pH 7.4, and 1 mM EDTA. The homogenate was centrifuged at 800 g for 10 min. Supernatant was filtered and centrifuged at 12,000 g for 15 min. Pellet was resuspended in buffer B containing 0.25 M sucrose, 75 mM mannitol, 10 mM TRIS HCl pH 7.4, 1 mM EDTA. The final supernatant was centrifuged at 12,000 g for 15 min and the mitochondrial pellet resuspended in Buffer B. 20 μg/ml ampicillin and protease inhibitor cocktail 50 μg/ml 5-fluorouracil were present throughout.

MIT was used just prepared for the evaluation of oxygen consumption and ATP synthesis, because it was observed that the OxPhos assay in mitochondria was extremely sensitive to the conditions of the sample: mitochondria can only synthesize ATP and consume O_2_ if they are assayed immediately after isolation, without prior freezing. After freezing the get uncoupled and do not synthesize ATP.

### Myelin isolation

Myelin was isolated from bovine forebrain according to the ‘‘floating up’’ modification (Haley *et al*. 1981) of the sucrose gradient method by Norton and Poduslo (Norton and Poduslo 1973). 0.5 g of each sample were homogenized with a Potter-Elvehjem homogenizer in 1.5 ml of 0.32 M sucrose in 2 mM EGTA. To avoid possible functional mitochondrial contamination, Cyclosporin A (Crompton *et al*. 1988) and 2-aminoethoxydiphenyl borate (Chinopoulos *et al*. 2003), inhibitors of the mitochondrial permeability transition pore opening, were not used during the preparations (Ravera *et al*. 2009). Homogenates were layered on top of 1 ml of 0.85 M sucrose in 2 mM EGTA and centrifuged at 75,000 g for 30 min in a Beckman FW-27 rotor (Beckman, Fullerton, CA, USA). This step was repeat twice. The pellet containing mitochondria, nuclei and cell debris was discarded. The sample at the interface between the two sucrose solutions, i.e. crude myelin (plasma membranes and myelin sheath) was retained, homogenized in water to a final volume of 1.5 ml, to obtain an hypo-osmotic shock, then centrifuged at 12,000 g for 10 min. Pellet was dispersed again in 1.5 ml of ultrapure water then centrifuged at 12,000 g for 10 min. Supernatant was discarded, and the densely packed pellet was dispersed in water and centrifuged at 75,000 g for 15 min. The final isolated myelin (IM) pellet was collected and stored at −80 °C. 20 μg/ml ampicillin and protease inhibitor cocktail 50 μg/ml 5-fluorouracil were present throughout.

### Purified bovine rod OS disks preparations

Purified rod OS were prepared in dim red light from 20 bovine retinas by sucrose/Ficoll 400 continuous density gradient centrifugation according to Schnetkamp and Daemen (Schnetkamp and Daemen 1982) at 4°C, in the presence of protease inhibitor cocktail and Ampicillin (100 μg/ml). Bovine eyes, obtained from a local slaughterhouse, were enucleated within 1 h from the animal death, retinas were extracted by filling the eye semicup with Mammalian Ringer (MR), for 10 min. MR composition was: 0.157 M NaCl, 5 mM KCl, 7 mM Na_2_HPO_4_, 8 mM NaH_2_PO_4_, 0.5 mM MgCl_2_, 2 mM CaCl_2_ pH 6.9 plus protease inhibitor cocktail and 50 μg/ml Ampicillin), and letting the retina to float freely. Retinas were gently vortexed in 600 mM sucrose, 5% w/v Ficoll, 10 mM glucose, 10 mM ascorbic acid, 1 mM CaCl_2_ and 20 mM Tris-HCl at pH 7.4. The filtered solution was layered on a continuous sucrose/Ficoll 400 gradient and the tubes centrifuged at 100,000 × g for 1 h. Isolated intact OS were collected from the ‘non leaky’ OS band. To avoid possible functional mitochondrial contamination, Cyclosporin A (Crompton *et al*. 1988) and 2-aminoethoxydiphenyl borate (Chinopoulos *et al*. 2003), inhibitors of the mitochondrial permeability transition pore opening, were not used during the preparations.

### Oxygen consumption measurements

O_2_ consumption was assayed in IM, rod OS disk and MIT by a thermostatically controlled oxygraph apparatus equipped with an amperometric electrode (Unisense – Microrespiration, Unisense A/S, Denmark) (Ravera *et al*. 2013; Panfoli *et al*. 2009). For each sample, 50 μg of total protein were used. The samples were incubated in the respiration buffer composed of: 120 mM KCl, 2 mM MgCl_2_, 1 mM KH_2_PO_4_, 50 mM Tris HCl, pH 7.4 and 25 µg/ml Ampicillin. Different respiring substrates were employed. In particular, to stimulate the complexes I + III + IV pathway, 5 mM pyruvate + 2.5 mM malate or 0.7 mM NADH were added to the respiration buffer, while to induce the respiration trough the Complexes II, III and IV pathway, 20 mM succinate was used. To verify that the oxygen consumption really depended by the respiring complexes, 50 μM rotenone (Complex I inhibitor) or 20 µM antimycin A (Complex III inhibitor) was added.

### Bioluminescent luciferase ATP assay

To evaluate the basal ATP synthesis in IM, rod OS disk and MIT by the ATP synthase, samples were incubated for 10 min at 37 °C in: 100 mM Tris-HCl (pH 7.4), 100 mM KCl, 1 mM EGTA, 2.5 mM EDTA, 5 mM MgCl_2_, 0.2 mM P1,P5-Di(adenosine-5′) pentaphosphate, 0.6 mM ouabain, ampicillin (25 µg/ml), 5 mM KH_2_PO_4_ and a specific respiring substrate. In particular, 5 mM pyruvate + 2,5 mM malate or 0.7 mM NADH were added to activate the pathway composed by Complexes I, III and IV, while 20 mM succinate was employed to induce the pathway formed by Complexes II, III and IV. ATP synthesis started after the addition of 0.1 mM ADP, ad was monitored for 2 minutes, in a luminometer (Glomax 20/20, Promega) by the luciferin/luciferase chemiluminescent method. ATP standard solutions between 10^−9^ and 10^−7^ M were used for calibration. The same experiments were performed in the presence of different ATP concentration (10-100 µM) in the incubation medium. Higher concentrations were not tested, as the amount of synthesized ATP would not be detectable in a high ATP background.

In other cases, to verify whether the sample was able to store the proton gradient to synthetize ATP later, samples were incubated in the same medium described above, but centrifuged at 20,000g for 5 minutes and resuspended in an incubation medium without any respiring substrate before the addition of ADP to induce the ATP synthesis.

### Statistics

Each data is representative of five experiments (n= 5), running in triplicate. Each experiment represents a different sample preparation. Multiple comparisons were performed using the analysis of variance (one-way ANOVA) followed by Bonferroni post hoc test. Analyses were performed using SigmaStat (Systat Software, Inc., San Jose, CA, USA) software. A P-value minor of 0.05 was considered as significant (p<0.05).

## RESULTS

Figure 1 reports the O_2_ consumption (Panels A-C) and ATP synthesis (Panels D-F) in mitochondria-enriched fractions (MIT), IM and rod OS. All samples consume oxygen and synthetize energy in the presence of conventional substrates. However, MIT and rod OS reach the maximum respiratory rates in the presence of pyruvate + malate, while IM displays the highest respiration rate after succinate addition. The virtual abolishment of respiration and energy production after the addition of rotenone and antimycin A, specific inhibitors of Complex I and Complex III, indicates that the OxPhos metabolism is due to the ETC. However, a comparison of the amount of ATP produced by the three samples shows evident quantitative differences. For example, in the presence of pyruvate + malate, MIT produces about 22 nanomoles of ATP/min/mg, while IM and rod OS synthetized about 43 and 425 nanomoles of ATP/min/mg, respectively.

**Figure 1:**
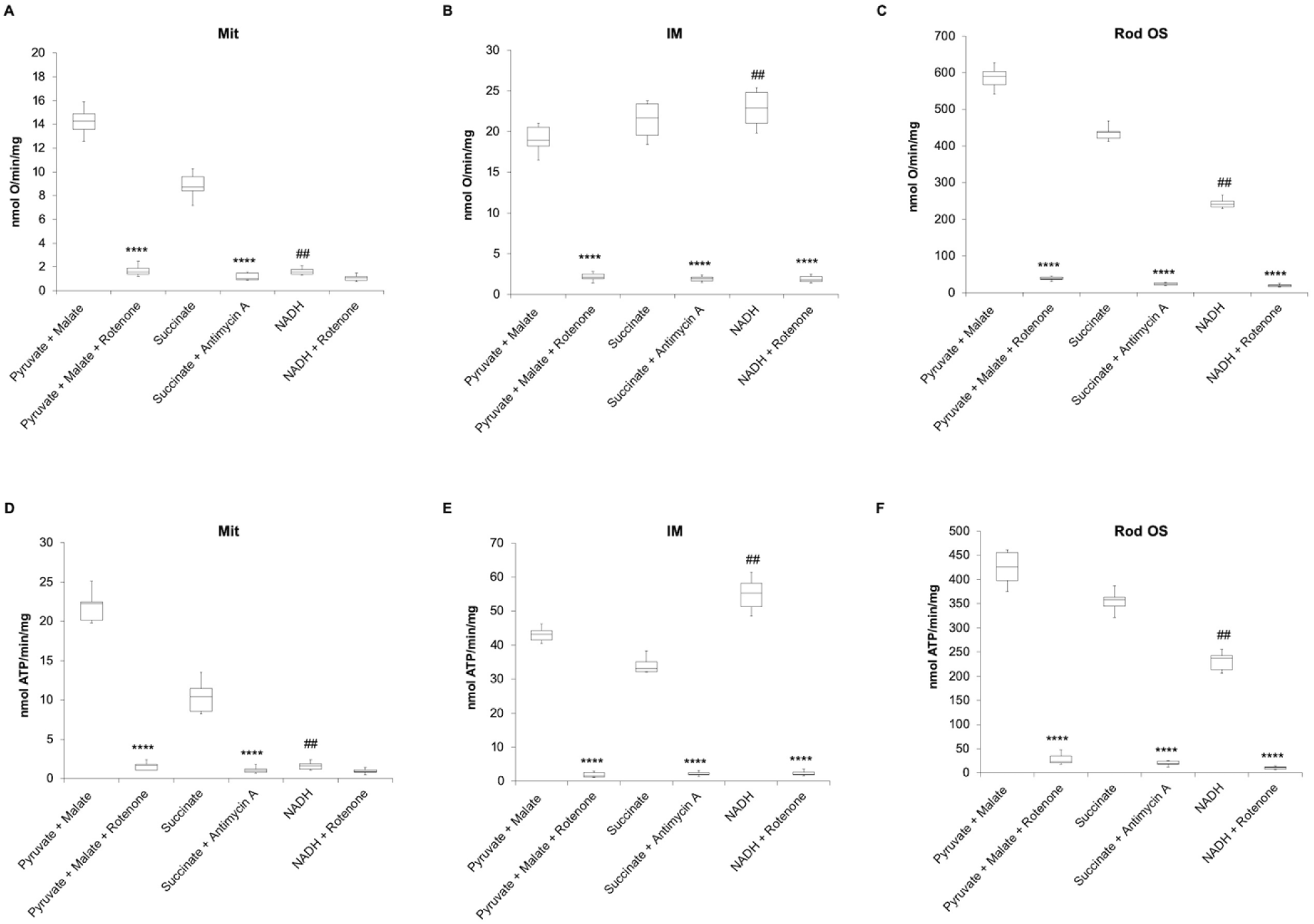
Oximetric and luminometric assays in mitochondrial enriched fraction, isolated myelin and rod outer segments. Panels A, B and C show the oxygen consumption performed by mitochondria enriched fraction (MIT), isolated myelin (IM) and rod outer segments (OS), respectively. Respiration was stimulated by the addition of pyruvate + malate, succinate or NADH, in the presence or in the absence of specific inhibitors (rotenone or antimycin A). O_2_ consumption is expressed as nmol O/min/mg. Panels D, E and F show the aerobic ATP synthesis in mitochondria enriched fraction (MIT), isolated myelin (IM) and rod outer segments (OS), respectively, induced by pyruvate + malate, succinate, or NADH, in the presence or in the absence of rotenone or antimycin A. Data are expressed as nmol ATP/min/mg. Each panel is representative of five experiments (n= 5), running in triplicate. **** indicates a significant difference of p<0.0001 between the absence or in the presence of the inhibitors of OxPhos activity. ## indicates a significant difference of p<0.01 between the oxygen consumption or ATP synthesis induced by pyruvate + malate or NADH.

Concerning the aerobic activity in the presence of NADH, IM show its maximum rate in the oxygen consumption and ATP synthesis and rod OS seems able to conduct an OxPhos with an unconventional substrate. Conversely, mitochondria do not display nor oxygen consumption nor ATP synthesis, confirming that NADH is not a suitable substrate for isolated mitochondria. As observed for the OxPhos activity in the presence by pyruvate + malate, rotenone determines decrement about 95% of oxygen respiration and energy production induced by NADH.

Testing the ATP synthesis induced by pyruvate + malate in the presence of different concentrations of ATP (0-100 µM) in the medium, we observed that the activity in MIT decreases proportional to the increment of ATP concentration, reaching an inhibition of 80% at 100 µM external ATP (Figure 2, Panel A). By contrast, the energy production in IM and rod OS seems not affected (Figure 2, Panels B and C, respectively).

**Figure 2:**
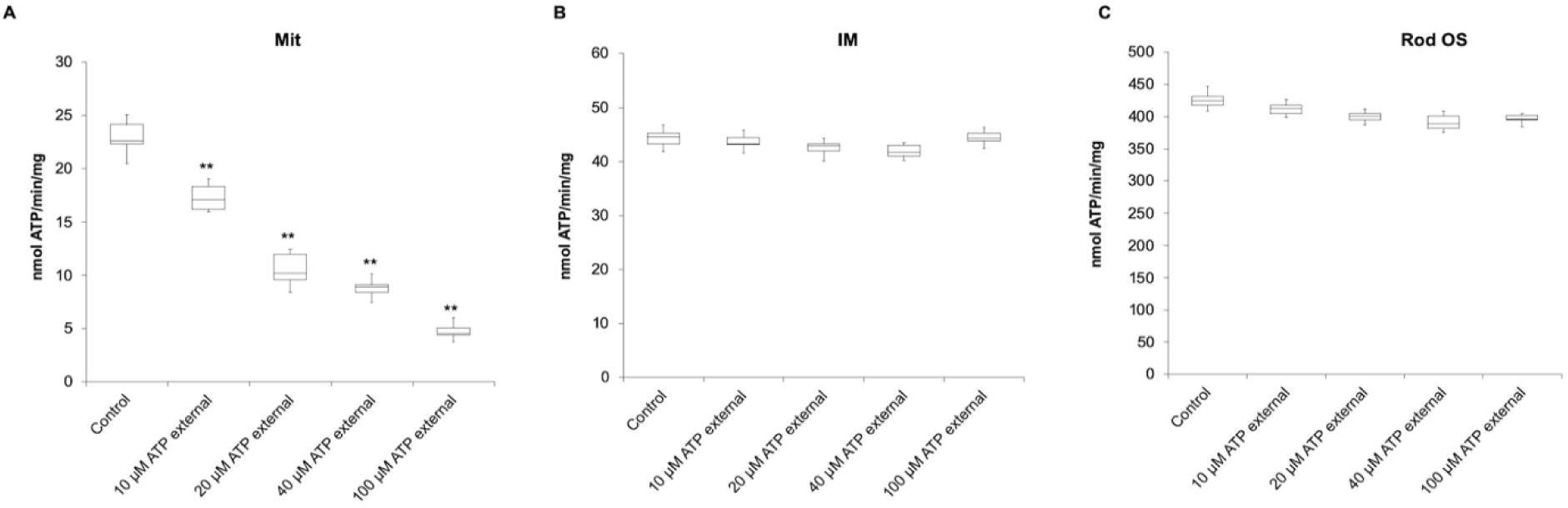
ATP synthesis assay in mitochondrial enriched fraction, isolated myelin and rod outer segments, in the presence of external ATP. Panels A, B and C report the ATP synthesis in mitochondria enriched fraction (MIT), isolated myelin (IM) and rod outer segments (OS), respectively, in the presence of different ATP concentration, ranging from 0 to 100 µM. ATP production is expressed as nmol ATP/min/mg. Each panel is representative of five experiments (n=5), running in triplicate. ** indicates a significant difference of p<0.01 between the sample in the absence of external ATP and the samples assayed after the addition of ATP in the medium.

In our previous work, we have hypothesized that ETC protons gradient may be accumulated into the membranes (Morelli *et al*.) (Morelli *et al*. 2011). Thus, to evaluate the protons accumulation, we have evaluated the ATP synthesis in the absence of respiratory substrates, in MIT, IM and rod OS pretreated from 10 to 30 minutes, with pyruvate + malate. In these conditions, MIT ATP synthesis appears negligible (Figure 3, Panel A), in comparison to that obtained in the presence of substrate during the assay and does not increase in parallel with the time of pre-incubation. Conversely, IM (Panel B) and rod OS (Panel C) synthesize ATP in the absence of respiring substrates in the assay medium, incrementing the production rate parallel with the incubation-time.

**Figure 3:**
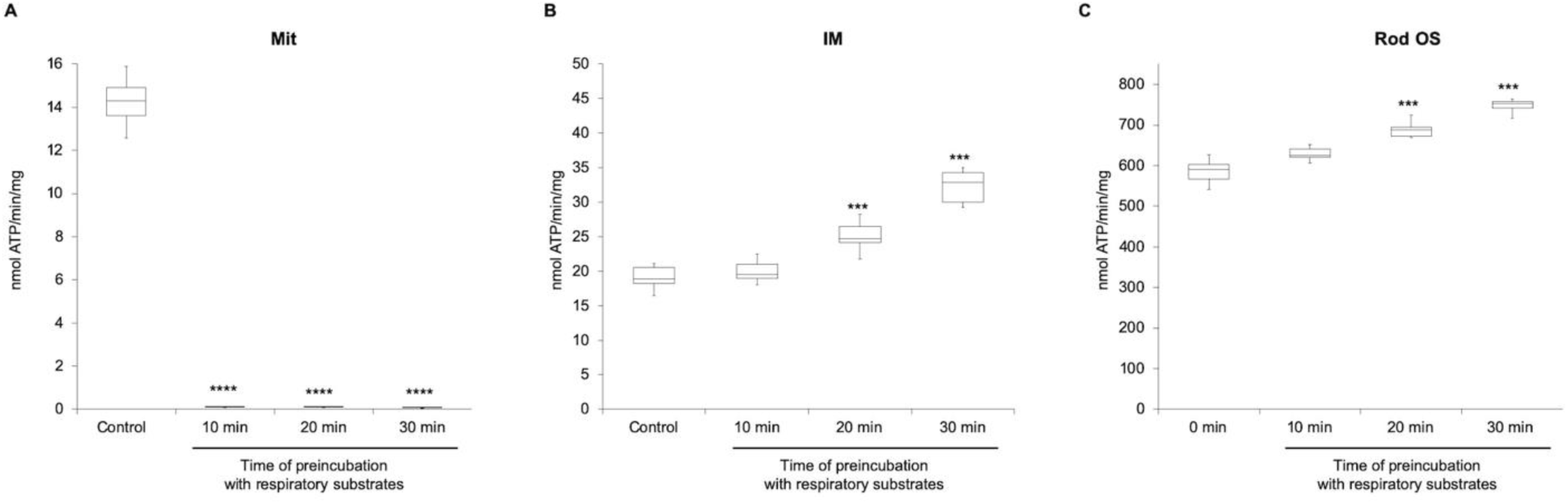
Evaluation of energy capacity accumulator by mitochondrial enriched fraction, isolated myelin and rod outer segments. Panels A, B, and C show a comparison of ATP synthesis assayed in the presence (control) or in the absence of respiring substrates, during the assay, in mitochondria enriched fraction (MIT), isolated myelin (IM) and rod outer segments (OS). In the case of substrates absence during the assay, the samples are previously incubated with 5 mM pyruvate + 2.5 mM malate, for increasing times (10-30 min). ATP synthesis is expressed as nmol ATP/min/mg. Each panel is representative of five experiments (n = 5), running in triplicate. *** indicate a significant difference of p<0.001 between the ATP synthesis assayed in the presence of respiratory substrate (control) and the ATP synthesis assayed in the preincubated samples before the analysis.

## DISCUSSION

In the present study, we compared the ability of MIT, IM, and rod OS disks to conduct the oxidative metabolism to investigate commonalities and differences of mitochondrial and extramitochondrial OxPhos metabolism.

Among the commonalities, all samples conduct the OxPhos metabolism after the induction of conventional substrates, pyruvate + malate or succinate, and are sensitive to the specific inhibitors of Complex I and Complex III. Interestingly, evaluating the rate of oxygen consumption and ATP synthesis, IM and rod OS seems more efficient in comparison to mitochondria. However, although the values obtained for mitochondrial ATP production by MIT are in line with those reported in literature (Eriksen *et al*. 2011) (Drew and Leeuwenburgh 2003) (Minet and Gaster 2010), it is important to note that both IM and rod OS maintain a structure similar to that observed *in vivo* (Ravera *et al*. 2009; Panfoli *et al*. 2009), while, during the isolation processes, mitochondria lose its reticulum organization, causing a decrement of OxPhos efficiency. In fact, it is note that isolated mitochondria are less performing with respect to those organized in a reticulum (Benard *et al*. 2007; Ravera *et al*. 2018a).

About the differences, only IM and rod OS conduct a complete OxPhos also in the presence of NADH. On the other hand, it is noted that NADH is considered a not suitable substrate for mitochondria *in vitro* because of the double membrane system and the absence of aspartate in the assay medium to activate the mitochondrial shuttles. Moreover, only the mitochondrial ATP synthesis is deeply affected by the presence of external ATP yield, while in IM and rod OS the activity of F_o_-F_1_ ATP synthase remains unaltered. This difference could be depending by the absence of Adenine nucleotide translocase (ANT) in IM and rod OS, as demonstrated in our previous works (Calzia *et al*. 2013a; Panfoli *et al*. 2009; Ravera *et al*. 2011; Ravera *et al*. 2013). In fact, ANT is involved in ATP transport, driven by the concentration gradient, from the mitochondrion to the cytosol (Liu and Chen 2013). Therefore, high external ATP concentration does not favor the mitochondrial ATP release. The ability of IM and rod OS to store the proton gradient to produce energy later represents another difference with mitochondria. This implies that a double membrane system is not necessary for the extramitochondrial OxPhos, to entrap the proton gradient. On the other hand, in our previous work, we have proposed a model in which protons from ETC would deliver inside the membrane, through specific transporters, such as cardiolipin and the phospholipid polar heads (Morelli *et al*.) (Morelli *et al*. 2019).

Interestingly, the IM ability to store the proton gradient could explain why the ATP production continues for about 13 min after the glucose deprivation (GD) (Trevisiol *et al*. 2017), although it has been proposed a contribute of astrocyte glycogen metabolism. Moreover, the ability to accumulate energy is in agreement with the differing vulnerability to ischemia of white matter in comparison to gray matter (Falcao *et al*. 2004). Similarly, the same capacity in rod OS to accumulate the proton gradient could account for the relative resistance, to a certain extent, to hypoxia of the photoreceptors, in spite of the retina is the highest oxygen-consuming tissue in the body (Braun *et al*. 1995).

Last, but not least the existence of differences between MIT, IM and rod OS OxPhos suggest that the extramitochondrial metabolism is not ascribed to a mitochondrial contamination during the sample preparation, as already demonstrated by western blot, immunofluorescence, and electron microscopy analyses in our previous works (Ravera *et al*. 2015; Panfoli *et al*. 2012; Ravera *et al*. 2013; Panfoli *et al*. 2009; Ravera *et al*. 2009; Ravera *et al*. 2011; Panfoli *et al*. 2011a; Calzia *et al*. 2013a). However, proteomic analyses show that the extramitochondrial OxPhos machinery is composed of the same subunits expressed in mitochondria, some of which codified by mitochondrial DNA (Panfoli *et al*. 2011b). This could be explained considering that the OxPhos machinery is organized in a “supercomplex” (Wittig *et al*. 2006; Lenaz and Genova 2009), which may be easily transferred from mitochondria to other membranous structures (Ravera *et al*. 2016; Bartolucci *et al*. 2015). On the other hand, mitochondria are dynamic organelle, able to fuse with other cellular compartments, such as the endoplasmic (de Brito and Scorrano 2010; Raturi and Simmen 2013), playing a pivotal role in the control of cell survival (Doghman-Bouguerra and Lalli 2019), calcium flux (Morciano *et al*. 2018), lipid metabolism (Flis and Daum 2013), and the entire energy metabolism (Rieusset 2018; Tubbs and Rieusset 2017; Lee and Min 2018; Marchi *et al*. 2014). A myelin that store energy opens up exciting prospects in the study of sleep, having been shown that sleep has been shown to lead to an increase in brain ATP (Dworak *et al*. 2010). Moreover exist clear links of sleep with the energy storage capacity (Caron and Stephenson 2010).

In conclusion, the OxPhos occurring in cellular compartments other than mitochondria represents a new vision of the ability of the cells to produce and storage energy, with exciting applications in many fields of biology.

## AUTHOR CONTRIBUTION

S.R., A.M. and I.P.: conceptualization; S.R.: data curation; S.R., M.B., and D.C.: investigation; A.M and I.P.: supervision; S.R., M.B.: writing - original draft; S.R., A.M. and I.P.: writing - review & editing.

